# DoubletDecon: Cell-State Aware Removal of Single-Cell RNA-Seq Doublets

**DOI:** 10.1101/364810

**Authors:** Erica A.K. DePasquale, Daniel J. Schnell, Íñigo Valiente-Alandí, Burns C. Blaxall, H. Leighton Grimes, Harinder Singh, Nathan Salomonis

## Abstract

Methods for single-cell RNA sequencing (scRNA-Seq) have greatly advanced in recent years. While droplet- and well-based methods have increased the capture frequency of cells for scRNA-Seq, these technologies readily produce technical artifacts, such as doublet-cell and multiplet-cell captures. Doublets occurring between distinct cell-types can appear as hybrid scRNA-Seq profiles, but do not have distinct transcriptomes from individual cell states. We introduce DoubletDecon, an approach that detects doublets with a combination of deconvolution analyses and the identification of unique cell-state gene expression. We demonstrate the ability of DoubletDecon to identify synthetic and cell-hashing cell singlets and doublets from scRNA-Seq datasets of varying cellular complexity. DoubletDecon is able to account for cell-cycle effects and is compatible with diverse species and unsupervised population detection algorithms (e.g., ICGS, Seurat). We believe this approach has the potential to become a standard quality control step for the accurate delineation of cell states.

## INTRODUCTION

The delineation of discrete transcriptomic cell states across organism development and disease will have profound impacts on biomedical research over the next several decades. Single-cell genomics provides a powerful means to derive and ultimately characterize novel cell states (Olsson et al., 2016; Velten et al., 2017; Villani et al., 2017; Yanez et al., 2017). While single cell profiling technologies continue to evolve at an astonishing pace, numerous challenges remain, including the separation of valid biological signatures from technical noise. A common source of confounding gene expression in single-cell RNA-Seq (scRNA-Seq) experiments is the occurrence of doublet or multiplet cell profiles, which result from the simultaneous capture of multiple cells in a single-well or droplet (Kang et al., 2018).

As demonstrated by cross-species mixing experiments, the frequency of single-cell doublets increases with the greater loading of cells for droplet-based scRNA-Seq platforms (Goldstein et al., 2017; Gong and Szustakowski, 2013; Macosko et al., 2015; Stoeckius et al., 2017). As a result, researchers are often advised to load fewer cells for these protocols to decrease the occurrence of doublets and hence limit the cellular depth afforded by these technologies. Beyond the mixing of individual cells, insufficient cellular dissociation will increase the frequency of cell aggregates which often result in hybrid expression profiles. The introduction of cell doublet profiles can significantly confound the experimental analysis and interpretation of scRNA-Seq data, in particular, the identification of novel cell states, developmental trajectories and biologically valid mixed-lineage progenitor states (Magella et al., 2017; Olsson et al., 2016). The identification of doublets from scRNA-Seq is further confounded by varying sparsity of the transcriptomic data, with as little as a few hundred unique molecular indexes (UMI) for a single-cell transcriptome, often resulting in poor correlation to comparable bulk RNA-Seq profiles (Kashima et al., 2018; Mantsoki et al., 2016). Although multiplet cell profiles should have a distinct global distribution of genes and UMI counts, these variables are insufficient to accurately predict which cells are doublets on their own (Stoeckius et al., 2017). Furthermore, other biological signals, such as cell-cycle effects, which are frequently variable in scRNA-Seq data (Scialdone et al., 2015), likely contribute to false positive doublet detection by simple synthetic doublet cell-profile analyses. While new emerging experimental methods, such as CITE-Seq and Cell-Hashing, enable the efficient identification of cell-multiplets, these approaches introduce additional experimental costs and cannot be applied to datasets analyzed without such protocols (Stoeckius et al., 2017).

Here we introduce a rigorous method for the identification of doublet-cell profiles from diverse scRNA-Seq platforms. Our method is able to account for common confounding biological signatures, such as cell-cycle effects and mixed lineage progenitor states that are not distinguished from real doublets by conventional *in silico* cell-mixing prediction approaches. Our approach applies a deconvolution method (nonnegative decomposition by quadratic programming), originally designed to estimate cell type proportions in bulk RNA-Seq data, to single-cell datasets to assess the underlying contribution of multiple concurrent cellular gene expression programs within a single-cell library. To prevent the inaccurate exclusion of transitional cell states associated with overlapping gene expression programs, we include additional steps to “rescue” these populations following initial doublet removal. We demonstrate the utility of our approach in a spectrum of developmental, disease and *a priori* identified doublet datasets.

## METHODS

### Algorithm Design

The software DoubletDecon has been developed as an open-source R package (https://github.com/EDePasquale/DoubletDecon) with a vignette on its use, algorithms and optional user-defined parameters. DoubletDecon identifies doublets through a three-step process: Remove, Re-cluster, Rescue (**Fig. 1**). Deconvolution profiles for single-cells are compared to real and synthetic doublet profiles, to maximize the representation distinct cell populations that are defined by few gene expression differences. Prior to these analyses, the software imports existing single-cell profiles, identifies distinct cell reference profiles and simulates doublet expression.

1. *<u>Data import and filtering</u>:* DoubletDecon takes input files from the standard outputs of the software ICGS (AltAnalyze) and Seurat, the latter of which are first processed through an accessory function within DoubletDecon (details of which can be found in the vignette). After processing, there is an optional cell-cycle gene cluster removal step to improve specificity of the algorithm.
2. *<u>Reference profile calculation:</u>* Gene expression medoids for each user-supplied cell cluster are calculated and correlations between the medoids are used to assess cluster similarity. The use of cluster medoids inherently excludes signal due to doublets within a cluster, though with highly sparse data it is advantageous to use a centroid. A binary correlation matrix is derived from the medoid correlations (*R*), which is termed the “blacklist”. The blacklist correlation threshold (*ρ’*) is user defined, with a default value of 1, with the resultant *ρ* used as a threshold for cluster similarity in the following formula:

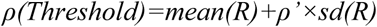 Higher values of *ρ’* will result in fewer blacklisted clusters. Markov clustering is used to define new clusters so that the similarity between clusters is minimized (Dongen, 2008) (**Fig. 1D**). This step is essential for heterotypic doublet removal, as homotypic doublets would not be expected to possess unique gene expression profiles. New medoids are created through this process and expression and the cluster identification is updated.
3. *<u>Synthetic doublet profile generation:</u>* To determine if the gene expression profile of a cell is more similar to an individual cell or doublet from two distinct cell-states, 30 synthetic doublets are generated for each pairwise combination of dissimilar clusters, with cells randomly sampled within each cluster and the resulting gene expression averaged.
4. *<u>Step 1 - Remove</u>*: The ‘Remove’ step of DoubletDecon uses deconvolution through quadratic programming with the R package ‘DeconRNASeq’ (Gong and Szustakowski, 2013). DeconRNASeq performs deconvolution on each cell expression profile, using all blacklisted cluster medoids as the references for deconvolution. The result will be a percentage estimate of the contribution of each reference cell-state (blacklisted cluster medoid) for queried cell (total always equal to 100%) (**Fig. 1E**). Each synthetic doublet centroid profile also undergoes deconvolution. For each cell in the dataset, the resulting deconvolution cell-profiles (DCP) are then compared to: 1) the centroid DCP for cells in each blacklisted cluster and 2) the centroid synthetic doublet DCP, using Pearson correlation. Individual cells with a DCP most similar to a synthetic doublet are subsequently predicted to be a doublet at this step.
5. *<u>Step 2 - Re-cluster</u>:* The cells that are removed as doublets are re-clustered in DoubletDecon’s ‘Re-cluster’ step. There are two user-defined options for re-clustering: 1) grouping of cells which have a common DCP (default) and 2) HOPACH clustering of the doublet gene expression profiles(Pollard and van der Laan, 2002). DCP group labels indicate the two highest correlated DeconRNASeq reference cell-types, alphabetically sorted (e.g., cluster-1 | cluster-2).
6. *<u>Step 3 - Rescue:</u>* In the ‘Rescue’ step of DoubletDecon, unique gene expression is assessed in the new doublet clusters via statistical enrichment (Welch’s t-test). Initial clustered doublets (e.g., B-cell | NK-cell) with at least one unique gene expressed relative to the original blacklisted clusters are re-assigned as singlets and reincorporated into the non-doublet expression matrix. For this analysis, DoubletDecon can evaluate all genes in the expression dataset or just those in the “marker” genes provided in the input files.
7. *<u>Downstream analyses:</u>* The results from DoubletDecon are immediately visualized (optional) via heatmap to evaluate the exclusion of visually distinct doublet gene expression signatures.

## RESULTS

To detect doublet profiles produced from very different cell types, as well as gradual cellular transitions, we developed a multi-step analysis strategy, that produces an initial set of putative doublets based on deconvolution analysis and then rescues erroneous doublet clusters that have unique gene expression (Methods, **Fig. 1A,B**). To test DoubletDecon in diverse use cases, we selected datasets with distinct biological and technical challenges for analysis (**Fig. 1C**). For this purpose, we used input data from the previously published ICGS and Seurat workflows, in which both preliminary unique cell states and cell-state associated genes sets were already defined or re-derived.

**Figure 1.**
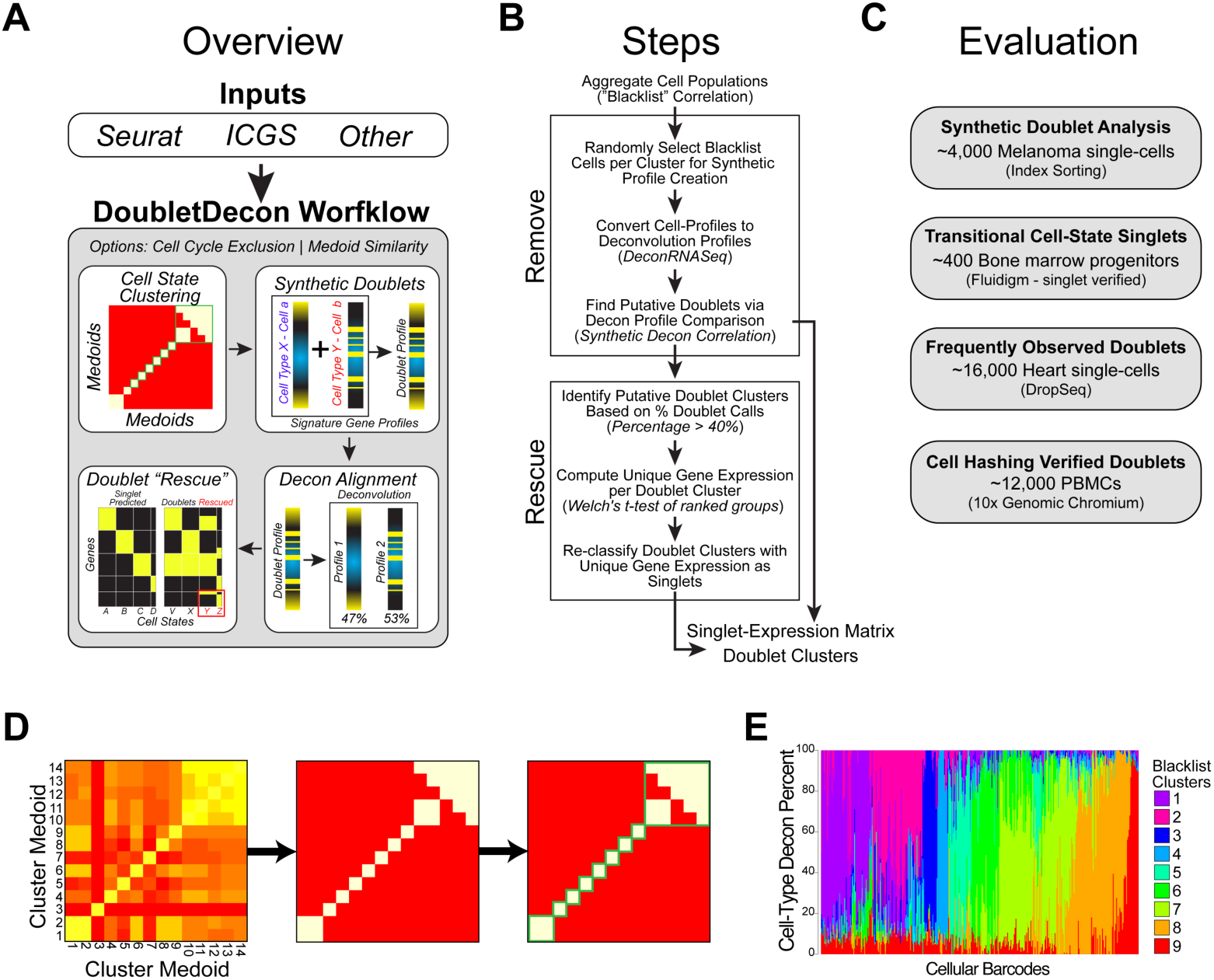
DoubletDecon workflow overview. A) Outline of the broad steps employed by this method, including cell state clustering with a “blacklist” of cluster medoids, synthetic doublet generation, deconvolution, and rescue through unique gene expression identification. B) Steps applied and output files from the remove and rescue portions of DoubletDecon. C) Datasets evaluated to assess DoubletDecon accuracy gene expression evidenced doublets. D) Cluster medoid blacklist determination. DoubletDecon identifies cell states which are closely related and thus are reasonable analogues for deconvolution analysis. Each medoid is calculated from the median gene expression of each separate cell-state for all algorithm selected cell-state marker genes (e.g., Seurat, ICGS). Initially, a medoid correlation matrix is created (left). Next, a threshold for medoid similarity is defined by the formula for *ρ* (outlined in the Methods), with the user-defined value of *ρ’* used to set the level of similarity required for a medoid to be considered correlated (middle). Finally, this new binary correlation matrix is visualized with a heatmap and Markov clustering is used to “square-off” the blacklist and determine which sets of clusters should not be considered for multiplet detection (right). E) The frequency of cell-state deconvolution profiles is shown for a dataset without doublets (microscopy validated, (Olsson et al., 2016)). Each column represents a different cell, in which each color indicates the percentage contribution of a reference cell-type (blacklisted) for that cell. Note, the majority are predicted to be composed principally of a single cell-type reference.

### Identification of Immune-Tissue Synthetic Doublets

As an initial test of DoubletDecon’s ability to detect doublets in a large and fairly deeply sequenced dataset, we used ICGS to identify cell populations and associated marker genes on over 4,000 cells from a previously described scRNA-Seq dataset of 19 melanoma tumors using the SMART-Seq2 protocol (Tirosh et al., 2016). From the provided processed expression data, we identified seven distinct immune (T-cell, B-cell, Macrophage) and non-immune cell-states (Melanocyte), in agreement with previously reported cell-populations (**Fig. 2A**). From this dataset we created 10 test sets containing synthetic doublets, which include the synthetic combination of immune and melanoma tumor cells. Briefly, the number of synthetic doublets created was 15 percent of the total number of cells, with cells from two different “blacklisted” clusters randomly chosen, and their gene expression averaged to create a synthetic doublet. These doublets were then inserted into the original ICGS-processed dataset using a recently developed cell matching and alignment algorithm called cellHarmony in the software AltAnalyze (AltAnalyze, 2017). This procedure was repeated 10 times to create datasets containing unique combinations of doublets. Each of the 10 datasets was tested with DoubletDecon 10 times to assess the variability in doublet calls between runs. Finally, a wide range of blacklist correlation thresholds (*ρ*’) were tested (ranging from 1 to 1.5 times the standard deviation above the mean correlation of the cluster correlation matrix, in increments of 0.05), as this is the primary variable in the function that is adjusted to account for differing similarities of clusters within the dataset (blacklisting).

**Figure 2.**
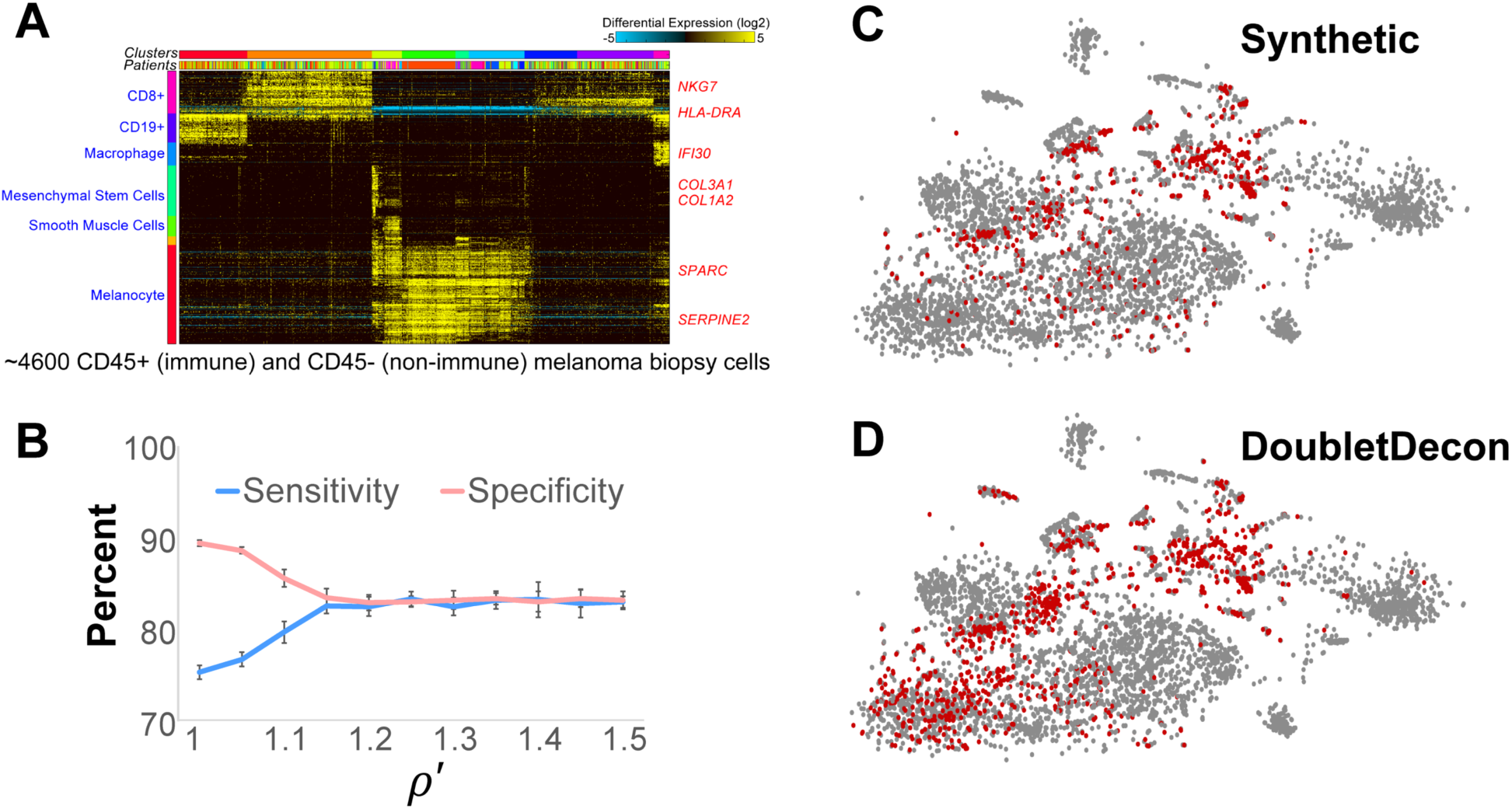
Results of doublet identification on synthetic doublets. A) ICGS analysis of 4,600 melanoma biopsy cells to identify gene and cell clusters, with associate gene-cluster cell-type enrichment predictions (blue text) and program assigned guide-genes (red text). B) Ten repeated trials on 10 separately generated sets of synthetic doublets resulted in a sensitivity and specificity estimates. C-D) Both synthetic doublets (C) and putative doublets (D) identified with DoubletDecon were visualized with t-SNE, with red dots indicating doublets and gray indicating non-doublets.

The results of these tests show an average sensitivity in detecting synthetic doublets of 83% and a specificity of 83% when the blacklist correlation threshold (*ρ’*) was greater that 1.2 (**Fig. 2B**). There was little variation in the resultant sensitivity and specificity across synthetic datasets and trials, which lends confidence to the reproducibility of these results (average standard deviation: sensitivity = 1.03%, specificity = 0.79%). Visualization of the predicted doublets and known synthetic doublets using t-Distributed Stochastic Neighbor Embedding (t-SNE) identified a similar distribution of synthetic and DoubletDecon-identified doublets (**Fig. 2C-D**). Notably, endogenous doublets in the original experiment likely exist but cannot be definitively verified.

### Rescue of Transitional Cell States Predicted as Doublets

Developmental and progenitor cell specification hierarchies inherently contain cells with transitioning gene expression profiles as well as mixed-lineage cell states as previously demonstrated (Magella et al., 2017; Olsson et al., 2016). Any doublet detection method will likely predict cells and cell-states associating with such transitions as doublets. In some cases, such predicted doublets will be defined by unique gene expression. Using a recently published hematopoietic bone marrow progenitor dataset of 383 cells with high-confidence assigned cell-types and singlet-restricted profiles (validated via cell capture imaging), we assessed the false-positive doublet call rate in these data. From this testing we find the maximum specificity to be 72% in the initial doublet detection step (“Remove”) but increased to 85% when unique gene expression was considered (“Rescue”) (**Table S1**). When comparing the number of cells called as putative doublets after step 1 (“Remove”) to those called as doublets after the final step (“Rescue”), up to 48% of the false positives are rescued and accurately identified as non-doublets. Hence, these results indicate that rescue of doublets is necessary to further reduce false positives (**Fig. 3A**). Further examination found that the proportion of cells that were recovered varied in different cell states, suggesting that similarity in the cell-state transcriptional profiles is a major contributor of doublet prediction (**Fig. 3B-C**). While ~40% monocytic progenitors are inaccurately predicted to be doublets, we note that recently cells in this cluster were determined to be either CMP derived and GMP derived, with subtle differences in their transcriptional profiles (Yanez et al., 2017). It is likely that these and other predicted doublet cell groups contain rare intermediate cells in cell populations that differ slightly in their gene expression but do not contain unique gene expression themselves.

**Figure 3.**
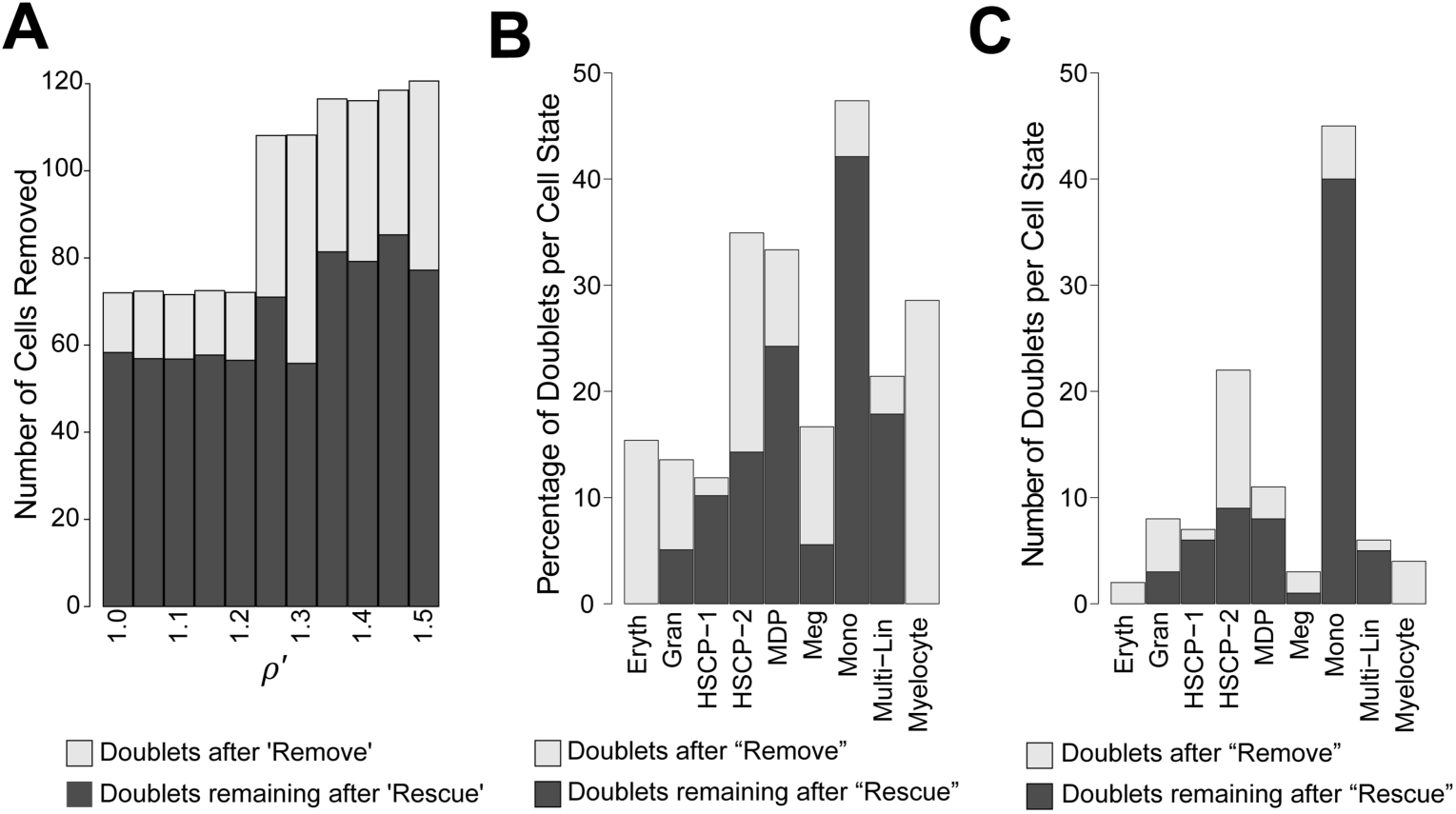
Results of doublet identification on verified hematopoietic progenitor singlets. A) DoubletDecon was run over a range of *ρ’* from 1 to 1.5 on Fluidigm-generated hematopoietic progenitor cells that have been visually inspected to confirm only singlets were present. The total height of the bars indicates the total number of cells that were identified as doublets after the “Remove” step of DoubletDecon across all values of *ρ’*, while the dark grey portions are those identified as doublets at the end of DoubletDecon over 10 independent trials. The difference in bar heights between the remove and rescue steps represent the number of cells that were “Rescued” by unique gene expression. B-C) These results are reported for a single representative trial for a single *ρ’* (1.3) for each previously annotated bone marrow progenitor population (Olsson et al., 2016). Both the number of cells (B) and percentage of cells per cell state (C) removed at each step are shown.

### Qualitative Assessment of Doublet Removal

In very large scRNA-Seq datasets (>10,000 cells), we expect hundreds to thousands of doublets to occur as separate distinct clusters (5-15%). ICGS analysis of >16,000 mouse heart cells collected through DropSeq (unpublished) identified 7 major cell populations, with visually identified doublet clusters and confounding cell-cycle effects (**Fig. 4A**). DoubletDecon predicted multiplet profiles show clear patterns of hybrid expression, including independent clusters with evidence of mixed cell-type expression (**Fig. 4A-C**). Although 20% of cells called as doublets in the “Remove” step of our analysis, more than twice what we would typically expect, the final results reported 11% of cells as doublets. These analyses were performed with cell-cycle cluster removal. Notably, without cell-cycle effects excluded, we found 26% doublets after the Remove step, however, DoubletDecon identifies 12% of these cells as doublets following the Rescue step. While most predicted doublets were interspersed throughout the major cell-type clusters, we note that nearly 50% of the cells in the last original cluster were identified as doublets, suggesting that this cluster is primarily composed of true-positive doublets. When this cluster was examined further, we discovered gene expression indicative of endothelial and fibroblast cells, which is a profile that we would not expect to normally occur, suggesting that all cells in the cluster are likely doublets.

**Figure 4.**
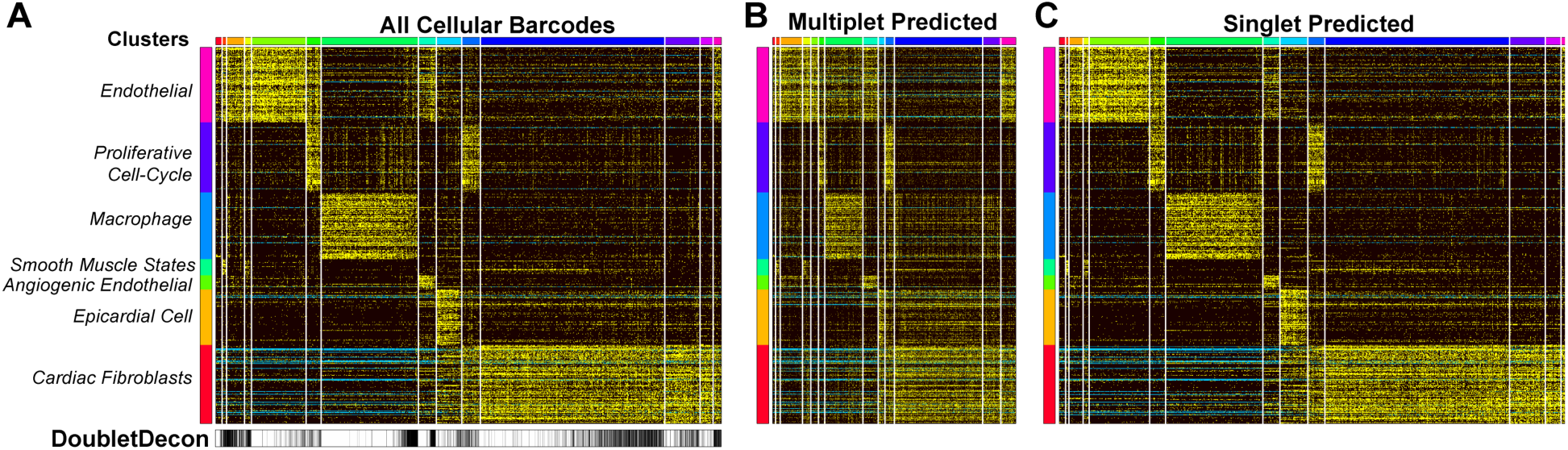
Visualization of DoubletDecon predicted multiplets in over 16,000 cells. A) Unsupervised clustering with ICGS was performed on approximately 16,000 heart cells collected through Drop-Seq and visualized in AltAnalyze, with potentially confounding gene expression clusters (e.g. cell-cycle effects) identified on the left. B-C) DoubletDecon predicted multiplets (B) and singlets (C) were also visualized in AltAnalyze. Input ICGS expression file is available from: https://www.synapse.org/#!Synapse:syn13849349/files/.

### Identification of Experimentally Verified Doublets from PBMC

As a positive test-case for known multiplet cell profiles, we analyzed a recently described human peripheral blood mononuclear cell (PBMC) dataset in which cell multiplets were experimentally defined based on a cell hashing barcoding strategy (Stoeckius et al. 2018) (Stoeckius et al., 2017). To obtain frequent doublets cell profiles, Stoeckius et al. overloaded a 10x Genomic Chromium port with hundreds of thousands of cells. A low gene expression cutoff was further applied by the authors to examine ultra-shallow single-cell profiles with as little as 200 unique molecular indexes. Using the same analysis workflow in the software Seurat, we selected the top 12,000 most highly expressed cells, only ~3,000 of which possessed over 500 genes expressed per cell. We predicted multiplets to be those cellular barcodes with >20% of the total hashing reads assigned to more than one hashing barcode. Visualization of these results by t-SNE in Seurat shows that hashing-based multiplets typically localize to the boundaries of distinct Seurat predicted clusters, though many are interspersed (likely homotypic doublets) (**Fig. 5A,B**). DoubletDecon, using centroids rather than medoids, to deal with increased data sparsity, identified up to 47% of Hashing annotated multiplets with 70% specificity (**Fig. 5C**). While generally low, these predictions were comparable to a recent alternative doublet detection algorithm called DoubletFinder (40% sensitivity and 93% specificity on the same Seurat-processed data) (McGinnis et al., 2018). The low sensitivity to detect such doublets is not surprising, given that many homotypic doublets are expected from the major cell populations detected by the unsupervised analysis.

**Figure 5.**
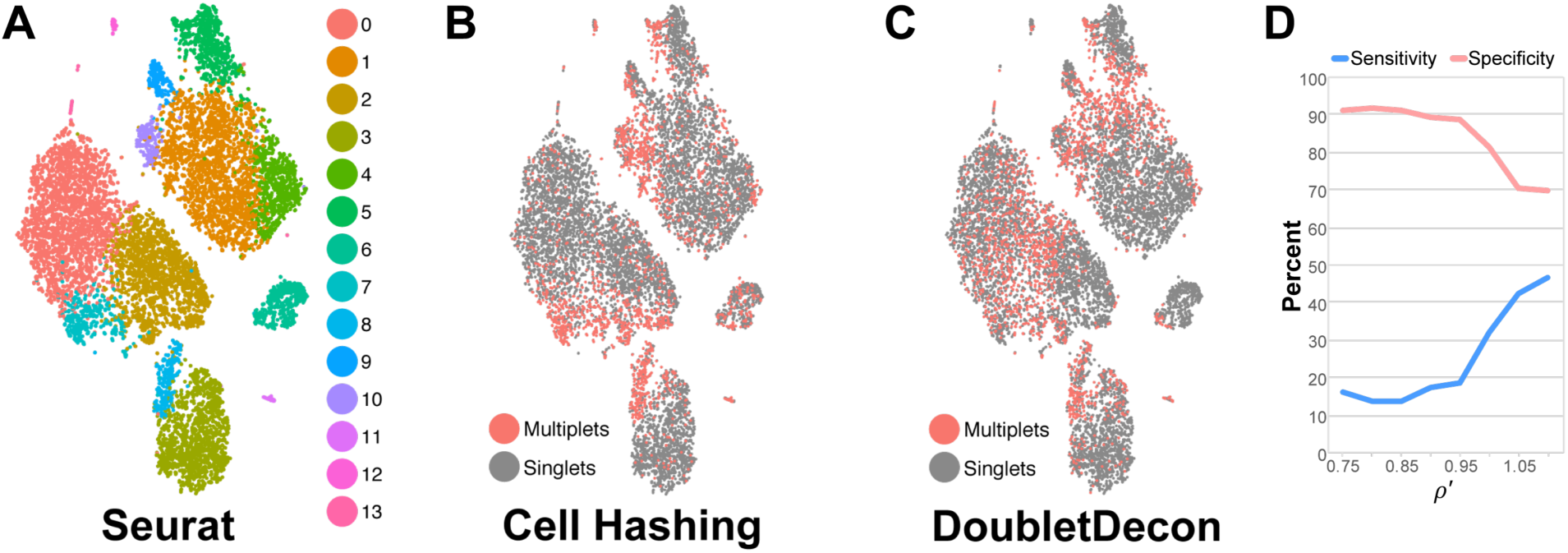
Performance of DoubletDecon on hashing-validated PBMC doublets. A) t-SNE plot of *de novo* clusters obtained from Seurat, performed on 12,000 PMBCs (after minimum gene expression and UMI filtering were applied, following cell-line associated cell profile exclusion as described by Stoeckius et al.). B) Cell hashing defined doublets are marked in red with known singlets in gray C). DoubletDecon predicted doublets (red) and singlets (gray) were visualized on the same t-SNE plot. D) Performance of DoubletDecon on cell-hashing barcodes run over the range of blacklist modifying *ρ’* values (0.75 to 1.1, default=1).

## DISCUSSION

As the number and cellular depth of single-cell datasets increases, standardized multiplet discovery workflows are necessary to remove technical cellular artifacts that can confound the identification of valid biological cell states. DoubletDecon accomplishes this goal by taking advantage of existing robust unsupervised population detection approaches, such as Seurat and ICGS, to effectively model doublet gene expression profiles. Our approach is applicable to both large and small datasets with both discrete cell populations or gradual cellular transitions, by automatically grouping correlated cells states without merging (blacklisted clusters). Given that valid hybrid transcriptomic states exist throughout development, such as transitional cell states and bi-potential intermediates, DoubletDecon includes specialized methods to “rescue” preliminarily “removed” cells and cell-state clusters that include unique gene expression patterns. As demonstrated here, this method significantly reduces the number of doublets which are known false positives and results in doublet frequencies similar to those expected with increased cell loading.

Although the initial version of DoubletDecon is able to effectively identify and exclude a high proportion of doublet gene expression profiles, we believe that this basic approach can be further expanded to identify additional unwanted and desired sources of variation. False negatives with this approach currently include multiplets of more than two cells, as well as doublets of highly similar cell states. We aim to enable the discovery of such multiplets in the future which should be readily identifiable using our existing deconvolution-based strategy. Minor variation in the results does occur due the random selection of cells for real and synthetic reference calculation. False positives currently include cells with intermediate lineage profiles that collectively do not result in unique gene expression. We believe this particular challenge can be overcome by improving our existing methods for unique population-associated gene expression or similarity of those profiles to singlets as additional selection steps, which remains an active area of research. In the analyses presented here, existing multi-lineage progenitors were not frequently classified as doublets and contained unique gene expression that resulted in the rescue of initially predicted doublet cells (Olsson et al., 2016). DoubletDecon further attempts to improve doublet removal through the exclusion of cell-cycle gene expression associated clusters. We anticipate that inclusion of custom or known confounding gene-sets will provide the flexibility to address additional known undesired effects. Ultimately, additional optimization and improvement of these methods will enable greater precision to characterize cells from samples with frequent doublets from diverse single-cell platforms and studies.

## ACKNOWLEDGMENTS

We acknowledge the assistance of the Cincinnati Children’s Hospital Medical Center (CCHMC) Single-Cell Genomics Core and DNA Sequencing Core. We thank Phillip Dexheimer for helpful discussions. This work was partly funded by CCHMC Research Foundation and NIH R01HL122661 (H.L.G.).

## AUTHOR CONTRIBUTIONS

EAKD and NS conceived the research, implemented the algorithm, performed the data analyses and wrote the manuscript. DJS provided statistical expertise and advice related to the algorithm. IV and BCB provided the heart single-cell DropSeq data. HLG, BCB and HS provided critical advice and oversight related to algorithm design and evaluation.

